# Lithium partially rescues gene expression and enhancer activity from heterozygous knockout of *AKAP11* while inducing novel differential changes

**DOI:** 10.1101/2025.02.24.639480

**Authors:** Nargess Farhangdoost, Alessia Pietrantonio, Yumin Liu, Calwing Liao, Daniel Rochefort, Alain Pacis, Martin Alda, Patrick A. Dion, Boris Chaumette, Anouar Khayachi, Guy A. Rouleau

## Abstract

Bipolar disorder (BD) is a complex psychiatric condition usually requiring long-term treatment. Lithium (Li) remains the most effective mood stabilizer for BD, yet it benefits only a subset of patients, and its precise mechanism of action remains elusive. Exome sequencing has identified *AKAP11* (A-kinase anchoring protein 11) as a shared risk gene for BD and schizophrenia (SCZ). Given that both the AKAP11-Protein Kinase A (PKA) complex and Li target and inhibit Glycogen Synthase Kinase-3 beta (GSK3β), we hypothesize that Li may partially normalize the transcriptomic and/or epigenomic alterations observed in heterozygous *AKAP11*-knockout (Het-*AKAP11*-KO) iPSC-derived neurons. In this study, we employed genome-wide approaches to assess the effects of Li on the transcriptome and epigenome of human iPSC-derived Het-*AKAP11*-KO neuronal culture. We show that chronic Li treatment in this cellular model upregulates key pathways that were initially downregulated by Het-*AKAP11*-KO, several of which have also been reported as downregulated in synapses of BD and SCZ post-mortem brain tissues. Moreover, we demonstrated that Li treatment partially rescues certain transcriptomic alterations resulting from Het-*AKAP11*-KO, bringing them closer to the WT state. We suggest two possible mechanisms underlying these transcriptomic effects: (1) Li modulates histone H3K27ac levels at intergenic and intronic enhancers, influencing enhancer activity and transcription factor binding, and (2) Li enhances GSK3β serine 9 phosphorylation, impacting WNT/β-catenin signaling and downstream transcription. These findings underscore Li’s potential as a therapeutic agent for BD and SCZ patients carrying *AKAP11* loss-of-function variants or exhibiting similar pathway alterations to those observed in Het-*AKAP11*-KO models.

## Introduction

BD is a severe psychiatric disorder characterized by recurrent episodes of mania or hypomania and depression^1,2^. It affects about 2.4% of the global population and is linked to reduced functioning, cognitive impairment, and a diminished quality of life^3,4^. BD patients are at a high risk of suicide with about a 20-30-fold increase compared to the general population^5^. The mood stabilizer Li has been shown to significantly lower the risk of illness recurrences, as well as decrease the risk of suicide and overall mortality rates^6^. However, only about 30% of BD patients are fully responsive to Li^7^. BD is highly heritable (∼70-90%^8–10^) but it is also very heterogeneous and has considerable genetic overlap with other psychiatric disorders, such as SCZ^11,12^. The largest-to-date whole-exome sequencing collaborative project for BD has identified *AKAP11* as a rare-variant large-effect risk gene shared with SCZ, showing an excess of rare protein-truncating variants (PTVs) in *AKAP11* in BD and SCZ cases compared to controls (combined odds ratio [OR] = 7.06, p = 2.83 × 10E−9)^13^. The *AKAP11* gene is under evolutionary constraint (LOEUF = 0.32, pLI =1 on gnomAD v4.0) and encodes AKAP11 protein, a membrane-associated and vesicular anchoring protein that is highly expressed in brain tissue^14–16^. The AKAP11 protein is known to bind to the regulatory subunits of cAMP-dependent PKA, directing it to different substrates for phosphorylation^17^. Specifically, AKAP11 functions as a scaffold protein to facilitate the phosphorylation of GSK3β, a key enzyme of the Wnt/β-catenin pathway^18^, via PKA^15,19^. Notably, GSK3β is a target of Li--the gold standard treatment for stabilizing the mood in BD patients^20–24^.

Previously, our team investigated the impact of heterozygous *AKAP11* loss-of-function (LoF) mutations on the gene expression landscape and profiled the epigenomic modifications, including histone H3 Lysine 27 acetylation (H3K27ac), a hallmark of active enhancers^25^. H3K27ac is a key epigenetic modification of histone H3 that marks active enhancers and is associated with active transcription of target genes^26–29^. We found widespread transcriptional dysregulation of genes primarily involved in DNA-binding transcription factor activity, actin binding and cytoskeleton regulation, as well as cytokine receptor binding functions. We specifically highlighted the role of aberrant intergenic and intronic enhancer activity in driving these gene expression changes^25^. Furthermore, we highlighted the significant downregulation of pathways related to ribosome structure and function, a pathway also altered in synapse proteomics from BD and SCZ post-mortem brain tissues, as well as heterozygous *Akap11*-KO mice^25,30^. However, the impact of Li on such transcriptomic and epigenomic changes is not yet understood. Given the known role of AKAP11-PKA protein complex as well as Li in inhibiting GSK3β through increasing serine 9 (S9) phosphorylation levels^15,19,31,32^, we postulated that further treating the Het-*AKAP11*-KO human iPSC-derived neurons with Li might reverse some of the transcriptomic and/or epigenomic modifications previously observed in Het-*AKAP11*-KO compared to wild-type (WT) neurons. In fact, Palmer et al.^13^ reported that, out of 11 BD cases carrying *AKAP11* PTVs with available clinical data on Li response, 7 were good responders, while 4 were not^13^. Notably, they showed that the percentage of good responders to Li among *AKAP11* PTV carriers (63.6%) was slightly higher than the overall response rate in BD patients in their study (52%)^13^. Although the sample size is too small to draw firm conclusions regarding *AKAP11* PTV carriers’ response to Li, these findings further emphasize the importance of investigating the direct molecular impact of Li in Het-*AKAP11*-KO models. Such research is a critical step toward identifying effective therapeutic strategies for this genetically and clinically heterogeneous disorder.

In this study, using RNA sequencing (RNA-seq), H3K27ac chromatin immunoprecipitation followed by sequencing (ChIP-seq), we first investigate the direct transcriptomic and epigenomics effects of Li treatment in Het-*AKAP11*-KO human iPSC-derived neurons compared to the untreated state. Subsequently, we compare the gene expression and intergenic and intronic H3K27ac profile of Het-*AKAP11*-KO-Li-treated (Het-*AKAP11*-KO-Li) vs. WT-untreated (WT-UNT) with Het-*AKAP11*-KO-untreated (Het-*AKAP11*-KO-UNT) vs. WT-untreated (WT-UNT) to uncover the transcriptomics and epigenomic effects of Li, including any rescue or novel changes, relative to previous findings with untreated Het-*AKAP11*-KO vs. WT^25^. We show that chronic Li treatment of Het-*AKAP11*-KO iPSC-derived neurons results in a limited number of differentially expressed genes (DEGs), most notably the downregulation of the glutamate ionotropic receptor N-methyl-d-aspartate (NMDA) type subunit 2A (*GRIN2A*). This gene has been identified as a candidate in a genome-wide association study (GWAS) investigating the lithium response in BD. Moreover, it has been associated with BD in a GWAS^33^ and is also a high-confidence SCZ risk gene identified in the largest schizophrenia sequencing study (OR: 24.1, P = 7.18 x 10-06)^34^. We next show that Li not only rescues some of the transcriptomic consequences of the heterozygous knockout of *AKAP11* but also induces new DEGs previously absent in Het-*AKAP11*-KO vs. WT. Finally, we noted two potential mechanisms via which Li may rescue or induce those DEGs and impact the transcriptome of Het-*AKAP11*-KO vs. WT: 1) by modifying H3K27ac levels at intergenic and intronic enhancers, influencing enhancer activity and transcription factor binding strength compared to the WT system; 2) by increasing the GSK3β S9 phosphorylation, thereby impacting WNT/β-catenin signaling pathway and the downstream genes regulated by this pathway.

By systematically comparing these different conditions, we pave the way for unraveling the complex transcriptomic and epigenomic impacts of chronic Li treatment in the Het-*AKAP11*-KO human iPSC-derived neuronal model. This approach sheds light on the underlying biological mechanisms and potential therapeutic opportunities for *AKAP11* LoF variant carriers, and possibly other BD or SCZ cohorts with similarly disrupted pathways as seen in Het-*AKAP11*-KO models.

## Results

### The effect of Lithium on the gene expression profile of Heterozygous *AKAP11*-KO iPSC-derived neurons

After chronically treating the 3 Het-*AKAP11*-KO clones chronically with Li (1.5 mM, 11 days), we first examined the direct gene expression modifications resulting from Li treatment in the Het-*AKAP11*-KO model (Het-*AKAP11*-KO-Li vs. Het-*AKAP11*-KO-UNT). We profiled the differentially expressed genes (DEGs) at adjusted P-value (p.adj) < 0.05 and |log2FC| > 0.25 and detected 33 total DEGS: 28 downregulated genes and 5 upregulated genes (Fig. 1a, b; Supplementary Table S1). Using a Z-score hierarchical clustering heatmap of DEGs, we determined the distinct averaged gene expression patterns of Het-*AKAP11*-KO clones with and without Li treatment (Fig. 1c; Supplementary Fig. S1c). Standing out among the DEGs was downregulation of *GRIN2A*, which encodes the GluN2A subunit of NMDA receptors--glutamate-gated cation channels characterized by high calcium permeability^35^ (Fig. 1b, c). To reveal functional differences between the Li-treated and untreated Het-*AKAP11*-KO groups and detect statistically enriched gene sets in our transcriptomic data, we carried out gene set enrichment analysis (GSEA)^36^ using Gene Ontology (GO)^37,38^ molecular function database. The top 10 most significantly enriched molecular functions were all upregulated in Het-*AKAP11*-KO-Li compared to *AKPA11*-KO-UNT (Fig. 1d). These functions include those related to RNA binding, processing, and protein synthesis (RNA binding, catalytic activity acting on RNA, snRNA binding, ribonucleoprotein complex binding, sequence-specific DNA binding, transcription regulator activity), ribosome regulation (structural constituent of ribosome, including both ribosomal protein and mitochondrial ribosomal protein genes), and transcription regulation (transcription regulator activity, sequence-specific DNA binding; Fig. 1d, Supplementary Table S2). Interestingly, our group previously reported that several of these molecular functions upregulated following Li treatment in Het-*AKAP11*-KO neurons—including RNA binding, structural constituent of ribosome, sequence-specific DNA binding, and transcription regulator activity—were significantly downregulated (p.adj < 0.05) in Het-*AKAP11*-KO-UNT vs. WT-UNT gene expression GSEA^25^ (Supplementary Fig. S1d, Supplementary Table S3). Specifically, pathways related to the structural constituent of ribosome, sequence-specific DNA binding, and transcription regulator activity have been also previously reported to be significantly downregulated (p.adj < 0.1) in the GSEA of synapse proteomics from post-mortem brain samples of BD and SCZ patients, compared to controls^30^.

**Fig. 1.**
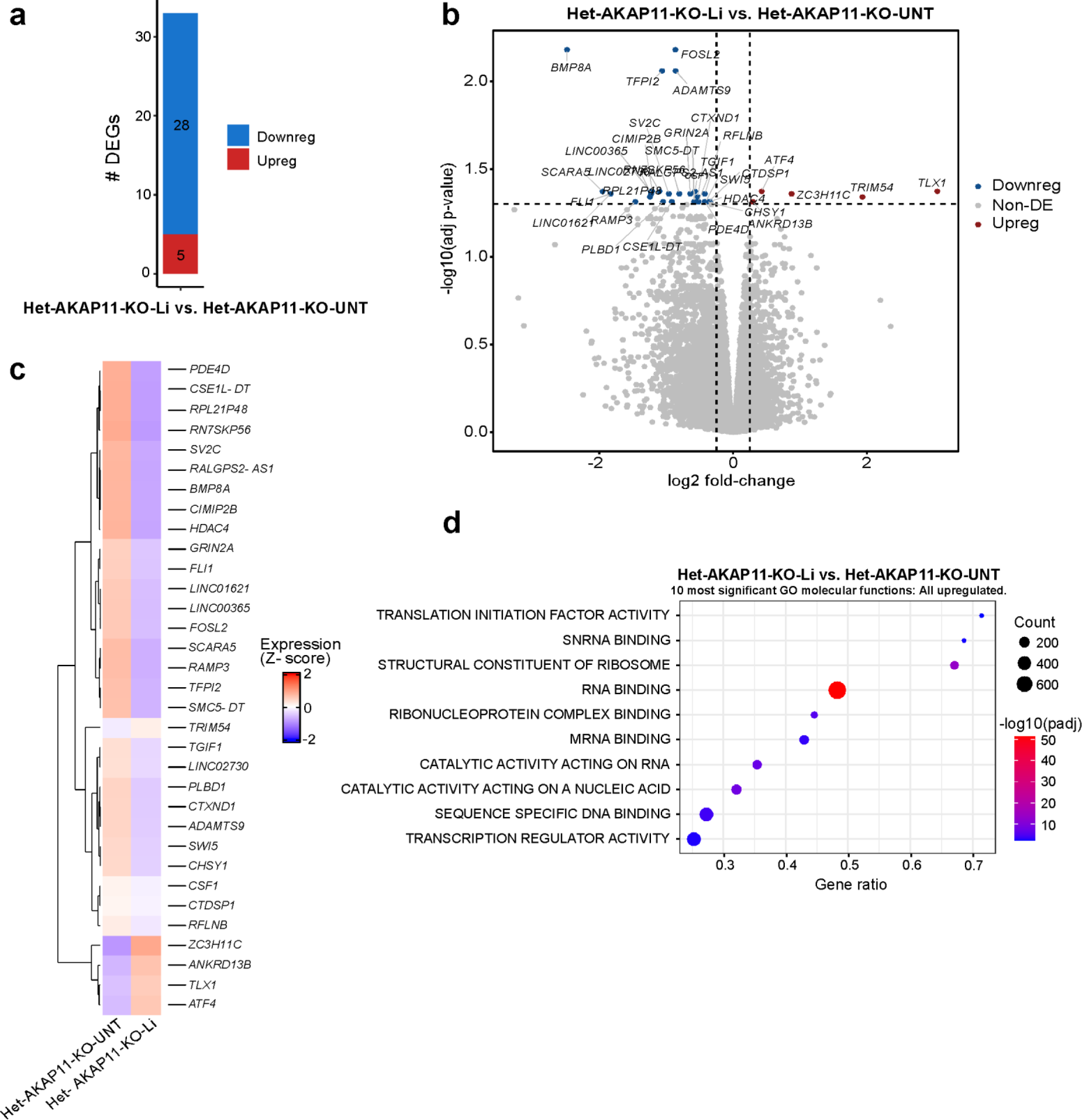
Li-dependent transcriptomic changes in Het-*AKAP11*-KO iPSC-derived neuronal model. (**a**) Number of up and downregulated DEGs in Het-*AKAP11*-KO-Li vs. Het-*AKAP11*-KO-UNT (P-value < 0.05 and |log2FC| > 0.25). blue: downregulated, red: upregulated. (**b**) Volcano plot of differentially expressed genes in Het-*AKAP11*-KO-Li vs. Het-*AKAP11*-KO-UNT (P-value < 0.05 and |log2FC| > 0.25). Red: upregulated, blue: downregulated, grey: nonDEG. (**c**) Hierarchical clustering heatmap of differentially expressed genes (rows) in Het-*AKAP11*-KO-Li clones vs. Het-*AKAP11*-KO-UNT reps (averaged). The expression levels across samples were standardized by the Z-Score method. Red shows the highest (2) and blue the lowest (-2) Z-score values. (**d**) GSEA^36,39^ showing the enrichment of the 10 most significant GO^37,38^ molecular function gene sets for the Het-*AKAP11*-KO-Li vs. Het-*AKAP11*-KO-UNT comparison; P.adj<0.05. For the full list refer to Supplementary Table S2. The size of the circles represents gene count (count of core enrichment genes); the colors represent the strength of -log (p-adj value), with red showing the most significant p-adj value, and gene ratio represents (count of core enrichment genes) / (count of pathway genes).

Li treatment resulted in more DEGs in the WT (WT-Li vs. WT-UNT, 87 DEGs, p.adj < 0.05 and |log2FC| > 0.25, Supplementary Fig. S1a, b; Supplementary Table S1) compared to the Het-*AKAP11*-KO neurons (Het-*AKAP11*-KO-Li vs. Het-*AKAP11*-KO-UNT, 33 DEGS). Interestingly however, Li impacted the gene expression profile of the WT neurons markedly differently with only 5 DEGs shared between the Li treatment within the WT and the KO models: *ADAMTS9*, *ATF4*, *BMP8A*, *LINC00365*, and *RFLNB* (Supplementary Table S1). These distinct responses to Li may be influenced by the baseline gene expression profiles of WT and Het-*AKAP11*-KO neurons, suggesting that different active pathways are directly targeted by Li. Nonetheless, we cannot interpret the results in relation to the quality of the therapeutic response to Li, as our investigation was conducted using iPSCs derived from a clinically unaffected individual.

### Lithium treatment of Heterozygous *AKAP11*-KO iPSC-derived neurons partially rescues KO-induced DEGs and induces novel DEGs

To identify the compensatory transcriptomic effects of chronic Li treatment in Het-*AKAP11*-KO neurons relative to WT-UNT, we first obtained DEGs from Het-*AKAP11*-KO-UNT vs. WT-UNT as well as Het-*AKAP11*-KO-Li vs. WT-UNT at p-adj < 0.05 and |log2FC| > 0.25 (Supplementary Fig. S2a, Supplementary Table S1). We observed an overall reduction in the number of DEGs, in both up-and downregulated directions, following chronic Li treatment of Het-*AKAP11*-KO neurons (Fig. 2a, Supplementary Fig. S2a, Supplementary Table S1). Li appeared to lessen the extent of differential gene expression that resulted from the heterozygous knockout of *AKAP11*, suggesting a partially compensating effect of Li on some of the Het-*AKAP11*-KO-induced gene expression changes. We then plotted a log2 fold change (Log2FC) heatmap showing the genes that were differentially expressed in Het-*AKAP11*-KO-UNT vs. WT-UNT but lost their DEG status upon Li treatment of the Het-*AKAP11*-KO (|log2FC difference|>0.6) hereafter referred to as Li-rescued DEGs (Fig. 2b, Supplementary Table S4). The heatmap shows that a relatively large portion of upregulated DEGs in the Het-*AKAP11*-KO vs. WT was rescued by treatment of Het-*AKAP11*-KO with Li (Fig. 2b). Among all the Li-rescued DEGs, we identified those encoding proteins involved in a variety of functions including DNA-binding transcription factor activity (*MEOX1, FOXC1, FOSL2, NKX3-2, IRX4, HOXD11, SHOX2, IRX3*), G protein-coupled receptor (GPCR) activity (*GAL, FZD10, RAMP3, SPHK1*), transmembrane transporter activity (*SLC7A8, ABCC3, SLC14A2, CHRND, GLRA1, RYR3*), excitatory extracellular ligand-gated ion channel activity (*CHRND, GLRA1*), growth factor binding (*FGFR3, DUSP1, FGFBP1, FGFBP2*), and oxidoreductase activity (*BCO1, HPD*) (Supplementary Fig. S2b). The most significantly enriched GO^37,38^ molecular function was glycosaminoglycan binding (*VTN, SMOC2, FGFBP1, HAPLN1, EGFLAM*; enrichment FDR=5.86E-02; Fig. 2d, Supplementary Fig. S2b). Subsequently, we showed that Li treatment of the Het-*AKAP11*-KO iPSC-derived neurons resulted in differential expression of a group of genes that were not differentially expressed in the Het-*AKAP11*-KO-UNT vs. WT-UNT comparison (|log2FC difference|> 0.6), hereafter referred to as Li-induced DEGs (Fig. 2c, Supplementary Table S4). Most of these genes went from being non-DEG in Het-*AKAP11*-KO-UNT vs WT-UNT to being differentially downregulated upon treatment of Het-*AKAP11*-KO with Li. ORA using the GO^37,38^ molecular function database (FDR < 0.1) revealed enrichment of extracellular matrix (ECM) structural constituent (*TNC, TFPI2, FMOD, LUM*), signaling receptor regulator activity (*FGF10, ANGPT4, WNT3A, NPPC, CALCB*), calcium channel activity (*TRPC7, OPRM1, TRPM2*), and metal ion transmembrane transporter activity (*TRPC7, SLC6A4, OPRM1, TRPM2, KCNJ16*) within these genes (Fig. 2e).

**Fig. 2.**
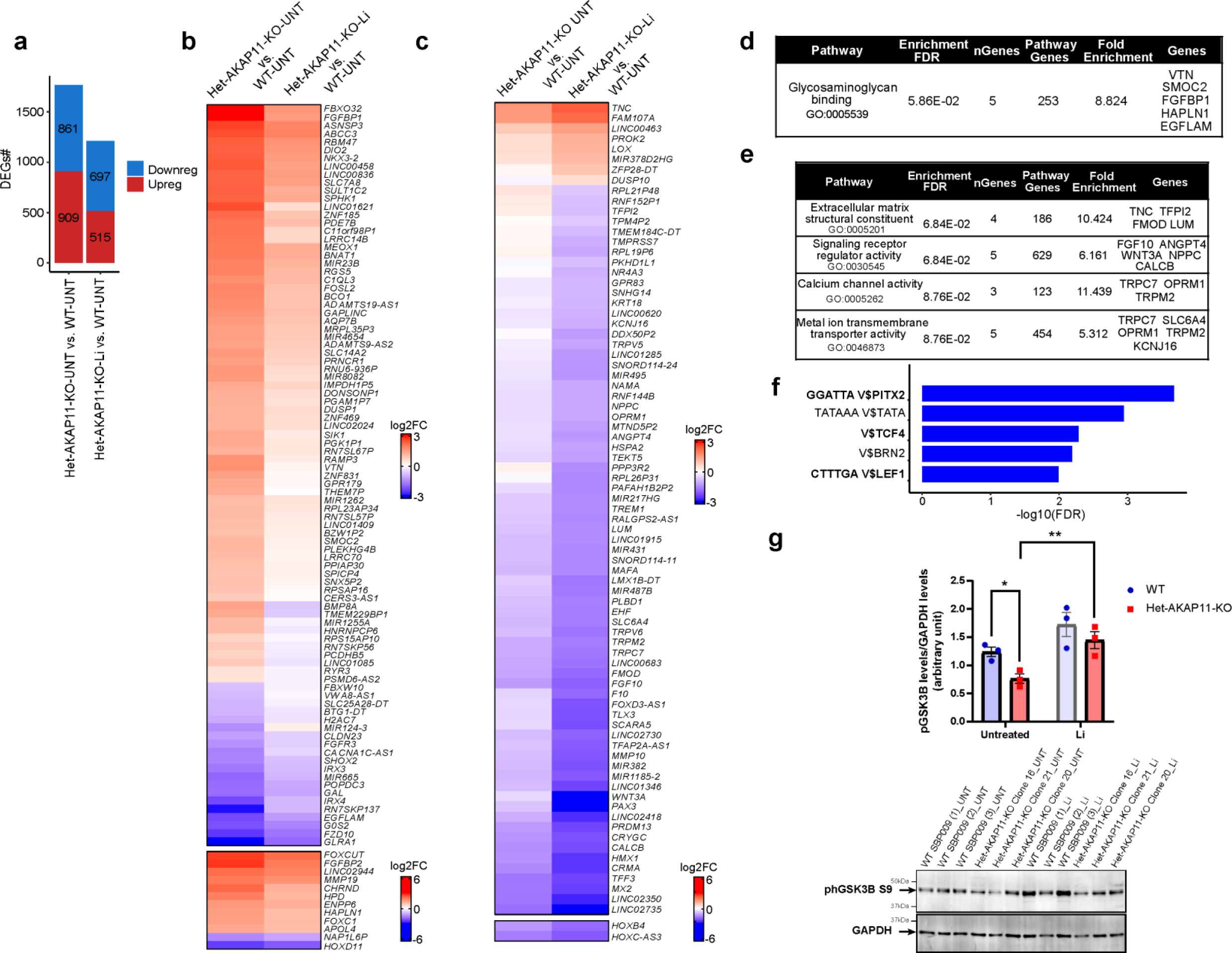
The differential gene expression rescuing and inducing effects of Li treatment in Het-*AKAP11*-KO relative to WT-UNT iPSC-derived neurons. (**a**) Number of up and downregulated DEGs in Het-*AKAP11*-KO-UNT vs. WT-UNT and Het-*AKAP11*-KO-Li vs. WT-UNT. P.adj < 0.05 and |log2FC| > 0.25. Red: upregulated, blue: downregulated. (**b**) Log2FC heatmap demonstrating the DEGs from Het-*AKAP11*-KO-UNT vs. WT-UNT that became nonDEGs (rescued) by Li treatment of the Het-*AKAP11*-KO (Het-*AKAP11*-KO-Li vs. WT-UNT). p-adj < 0.05 and |log2FC| > 0.25; DEG cutoff: P-adj < 0.05 and |log2FC| > 0.25. A minimum log2FC difference of |0.6| between the two sets of comparisons was set as a condition for [(Het-*AKAP11*-KO-Li vs. WT-UNT) – (Het-*AKAP11*-KO-UNT vs. WT-UNT)]. (**c**) Log2FC heatmap demonstrating genes that were nonDEGs in Het-*AKAP11*-KO-UNT vs. WT-UNT and became DEGs (induced) in Het-*AKAP11*-KO-Li vs. WT-UNT. p-adj < 0.05 and |log2FC| > 0.25; A minimum log2FC difference of |0.6| between the two sets of comparisons was set as a condition (Het-*AKAP11*-KO-Li vs. WT-UNT) – (Het-*AKAP11*-KO-UNT vs. WT-UNT). (**d, e**) Over-representation analysis (ORA) of rescued and newly induced DEGs, respectively, following Li treatment of Het-*AKAP11*-KO, using all genes from Figs. 2B and 2C heatmaps. GO^37,38^ Molecular function database using ShinyGO 0.77^40^; FDR < 0.1. The background genes used were all the 19,568 genes that were considered expressed (see Methods). Shown here are the top 10 pathways with redundancy removed. (**f**) Top 5 (based on adj p-value) Transcription Factor Target Genome Browser PWMs (FDR<0.05) enrichment results used to detect which binding sites were most significantly associated with the Li-responsive DEGs. Determined using ShinyGo 0.77^40^. For the full list of the top 20 motifs refer to Supplementary Table S5. (**g**) Bottom: Cropped Immunoblot membranes showing the levels of phGSK3β (S9) and GAPDH (loading control) on the same gel in the Het-*AKAP11*-KO clones and WT replicates with and without Li treatment. The bands indicated with the arrows were quantified. Full blots are presented in Supplementary Fig. S2c. Top: A two-way ANOVA followed by Fisher’s LSD post-hoc statistical test was conducted to determine the statistical significance of differences in phosphorylation levels of GSK3β among the Het-*AKAP11*-KO and WT groups. Data are represented as mean ± SEM. Statistical significance is indicated by NS (not significant), *p<0.05, **p<0.01, ***p<0.001.

It is important to emphasize the presence of the key WNT/β-catenin-pathway genes, such as *FZD10* and *WNT3A* (Fig. 2b, c), as well as the enrichment of β-catenin-regulated TCF4, LEF1, and PITX2 binding sites within the Li-responsive genes (Li-rescued and-induced DEGs). The latter was determined by ORA using the Transcription Factor Target Genome Browser Position Weight Matrix database (PWMs; enrichment FDR<0.05; Fig. 2f, Supplementary Table S5). This suggests that the dysregulation of GSK3β phosphorylation at the S9 residue—targeted by both AKAP11-PKA and Li—may play a role in Li-response-related transcriptomic modifications. These modifications include partial rescuing of gene expression changes caused by the heterozygous knockout of *AKAP11*, as well as the induction of novel gene expression changes that were previously absent. To test this hypothesis, we carried out immunoblotting of phospho-GSK3β (phGSK3β, S9) in all our samples, Li-treated and untreated, and quantified the bands (Fig. 2g, top and bottom; Supplementary Fig. S2c; Supplementary Table S6). We observed that heterozygous knockout of *AKAP11* led to a notable reduction (p=0.047) in the phosphorylation of GSK3β at Ser9 compared to WT neurons. Subsequent treatment with Li significantly increased Ser9 phosphorylation levels (p=0.009) in Het-*AKAP11*-KO neurons relative to their untreated counterparts, restoring levels toward, and even exceeding, those observed in the WT baseline (Fig. 2g, top).

### Impact of lithium on differential intergenic and intronic H3K27ac modifications resulting from heterozygous knockout of *AKAP11* relative to WT: implications for regulation of lithium-rescued and induced gene expression changes

We next aimed to determine whether modifications in the H3K27ac mark, a hallmark of active enhancers^27–29^, are involved in modulating gene expression changes resulting from chronic Li treatment in the Het-*AKAP11*-KO system. With this purpose in mind, we first profiled the intergenic and intronic H3K27ac differential peaks, which are the primary sites of enhancers previously dysregulated with the heterozygous knockout of *AKAP11*, in Het-*AKAP11*-KO-Li vs. Het-*AKAP11*-KO-UNT at p.adj < 0.05 and |log2FC| > 0.25 and detected 39 intergenic and 44 intronic differential peaks; Supplementary Fig. S3a, b; Supplementary Table S7). However, within the WT system (WT-Li vs. WT-UNT) there were no differential H3K27ac peaks with the same cutoffs of p.adj < 0.05 and |log2FC| > 0.25, highlighting that the H3K27ac modifications observed above are Het-*AKAP11*-KO-specific (Supplementary Table S8). We then associated the differential H3K27ac peaks from Het-*AKAP11*-KO-Li vs. Het-*AKAP11*-KO-UNT with the nearest TSS of known genes—within the distance of 100 kb from the TSS to the peak center—and obtained a total of 28 different genes associated with intergenic and intronic differential H3K27ac peaks and marked their gene expression status (Supplementary Table S7); however, there was no overlap between them and the 33 DEGs from gene expression analysis of Het-*AKAP11*-KO-Li vs. Het-*AKAP11*-KO-UNT at p.adj < 0.05 and |log2FC| > 0.25.

Next, we sought to investigate whether Li treatment rescues or further modifies the Het-*AKAP11*-KO-induced differential H3K27ac peaks in Het-*AKAP11*-KO iPSC-derived neurons relative to WT. We specifically aimed to detect modifications that aligned with Li-rescued or Li-induced DEGs, with respect to the direction of change (up/down) and the type of effect (induced/rescued). Like our gene expression data, we observed an overall reduction in the number of both differential intergenic and intronic H3K27ac peaks in Het-*AKAP11*-KO-Li vs. WT-UNT relative to Het-*AKAP11*-KO-UNT vs. WT-UNT, in both up and down directions (Fig. 3a, b, Supplementary Table S8). We specifically searched for the intergenic and intronic differential H3K27ac peaks in Het-*AKAP11*-KO vs. WT that lost their differential status (Li-rescued peaks) as well as intergenic and intronic differential H3K27ac peaks that newly appeared in Het-*AKAP11*-KO-Li vs. WT-UNT (Li-induced peaks; p.adj < 0.05 and |log2FC| > 0.25 for both, |log2FC difference| > 0.3; Supplementary Table S9). We then associated these Li-responsive peaks (both rescued and Li-induced peaks) with the nearest TSS of known genes within the distance of 100 kb from the TSS to the peak center. Subsequently, we integrated the H3K27ac with gene expression data and detected 8 intergenic peaks (associated with 8 genes) and 11 intronic peaks (associated with 9 genes) among the Li-rescued and Li-induced peaks that were overlapping our Li-rescued or Li-induced DEGs, respectively, with the same direction of change (Fig. 3c, d, Supplementary Table S9). In particular, one gene (*PLEKHG4B*) has both intergenic and intronic Li-rescued H3K27ac peaks. Although a few Li-induced DEGs were noted among these genes, most were Li-rescued DEGs (Fig. 3c, d). Among all these Li-induced and Li-rescued changes in enhancer activity that were linked to Li-induced and Li-rescued gene expression changes in the same direction, there were several ncRNA genes (including lncRNA, miRNA, and anti-sense RNA). In addition, we noted several coding genes playing roles in SLC-mediated transmembrane transport (*SLC7A8, SLC14A2*) and MAPK family signaling cascades (*FGFR3, DUSP1*), calcitonin-like ligand receptors (*RAMP3*) among others (Supplementary Table S10). Finally, we demonstrated the gene expression and H3K27ac averaged tracks of Li-treated and -untreated Het-*AKAP11*-KO clones, as well as untreated WT replicates, using the UCSC genome browser. The example tracks highlight the gene *RAMP3* which demonstrates an intergenic differentially elevated H3K27ac peak in Het-*AKAP11*-KO vs. WT, marking increased enhancer activity and differential binding, along with upregulation of gene expression, both of which are then lowered to a non-differential state in Het-*AKAP11*-KO-Li vs. WT-UNT, and thereby rescued (Fig. 3e).

**Fig. 3.**
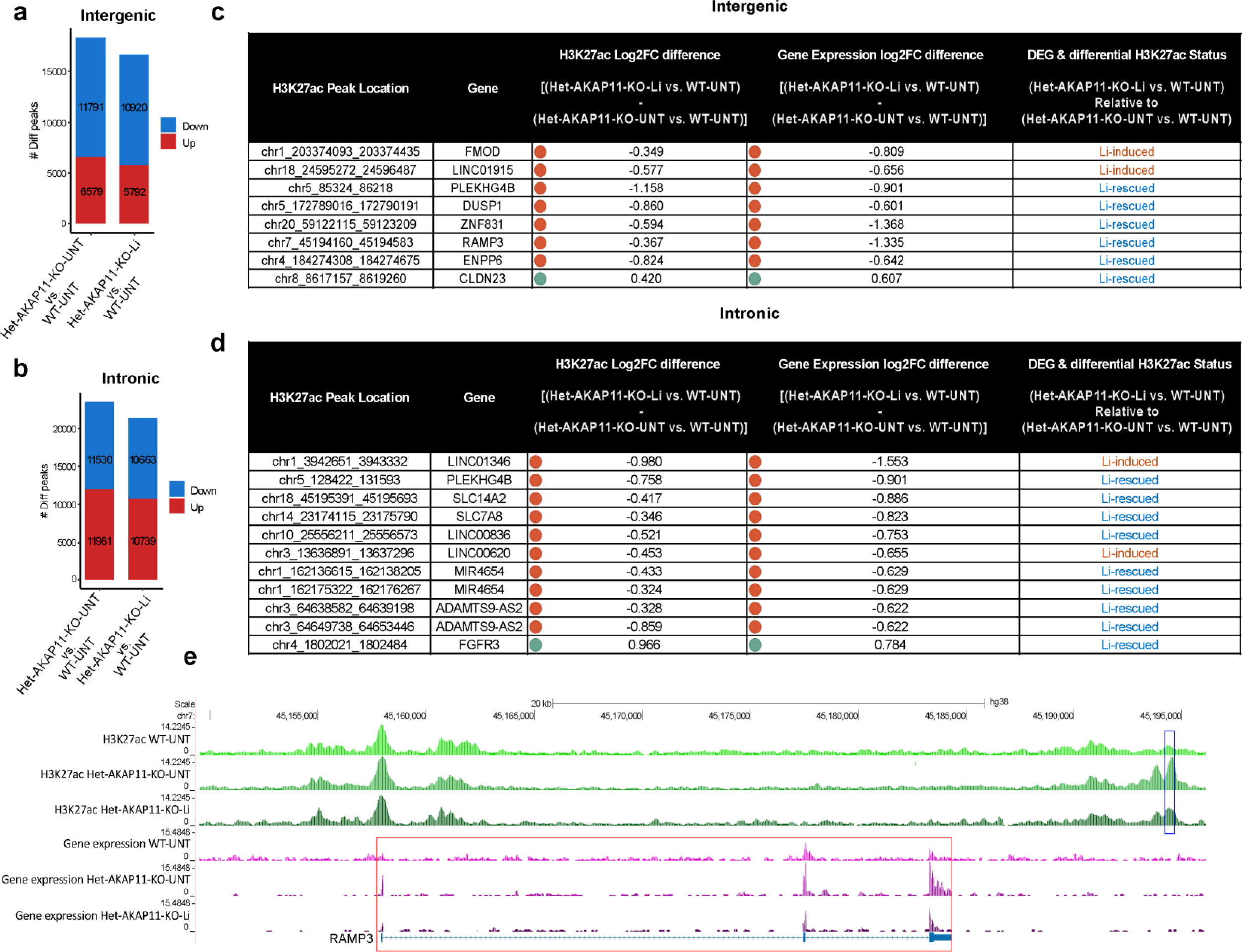
Li-rescued/induced H3K27ac peaks and their association with Li-rescued/induced gene expression changes: differential H3K27ac and gene expression modifications in Het-*AKAP11*-KO-Li vs. WT-UNT and Het-*AKAP11*-KO-UNT vs. WT-UNT. (**a, b**) Number of differentially elevated and decreased H3K27ac peaks in intergenic and intronic regions, respectively, in Het-*AKAP11*-KO-UNT vs. WT-UNT and Het-*AKAP11*-KO-Li vs. WT-UNT; p.adj < 0.05 and |log2FC| > 0.25. Red: differentially increased peaks, blue: differentially decreased peaks. (**c, d**) Tables showing Li-rescued/induced differential H3K27ac peaks that changed in the same direction as the Li-rescued/induced DEGs, respectively, following chronic Li treatment of Het-*AKAP11*-KO (Het-*AKAP11*-KO-Li vs. WT-UNT relative to Het-*AKAP11*-KO-UNT vs. WT-UNT). For both gene expression and H3K27ac: differentially marked (P.adj < 0.05 and |log2FC| > 0.25) in one comparison but not the other (i.e., in either Het-*AKAP11*-KO-UNT vs. WT-UNT or Het-*AKAP11*-KO-Li vs. WT-UNT); |log2FC difference| > 0.3 between the two sets of comparisons [(Het-*AKAP11*-KO-Li vs. WT-UNT) – (Het-*AKAP11*-KO-UNT vs. WT-UNT)] was also set as a condition. Red circle shows a negative value of Log2FC difference as a result of (Het-*AKAP11*-KO-Li vs. WT-UNT) – (Het-*AKAP11*-KO-UNT vs. WT-UNT); green circle shows a positive value of Log2FC difference as a result of (Het-*AKAP11*-KO-Li vs. WT-UNT) – (Het-*AKAP11*-KO-UNT vs. WT-UNT). Li-induced (non-differential-> differential) DEG/differential peaks are colored in orange and Li-rescued (differential->non-differential) are colored in blue. (**e**) UCSC Genome Browser H3K27ac and gene expression averaged tracks of Het-*AKAP11*-KO clones and WT replicates, Li-treated and untreated. Demonstrating *RAMP3*, as an example, with Li-rescued expression profile and an intergenic Li-rescued H3K27ac peak. Blue rectangles highlight enhancers with differential binding strength and red squares mark gene expression profiles of the genes.

These findings suggest that one mechanism by which Li rescues some of the DEGs resulting from the heterozygous knockout of *AKAP11* or induces new DEGs is by modulating intergenic and intronic H3K27ac levels in a concordant direction. This indicates that Li may play a role in modifying enhancer activity and transcription factor binding at intergenic and intronic regions, thereby rescuing or inducing gene expression patterns compared to the WT system.

## Discussion

BD is a highly heritable and heterogeneous psychiatric condition, and Li remains one of the most effective mood stabilizers for its treatment^6^. Studying the molecular effects of Li in Het-*AKAP11*-KO neuronal culture could contribute to refining treatment strategies and improving outcomes for this rare LoF risk gene associated with both BD and SCZ. Given the identification of *AKAP11* as a rare PTV large-effect risk gene shared between BD and SCZ^13^, and its involvement at the protein level in GSK3β regulation—a pathway influenced by Li^13,18^—in this study we: 1) used bulk RNA-seq data to first investigate how chronic Li treatment of Het-*AKAP11*-KO iPSC-derived neurons impacts the gene expression landscape within this model. Subsequently, we performed ChIP-seq of H3K27ac mark, a histone modification indicative of enhancer activity and transcription factor binding, to determine if changes in this histone mark are associated with the dysregulation of gene expression exerted by Li. 2) used bulk RNA-seq data to explore the Li-rescued and-induced differential gene expression by comparing Het-*AKAP11*-KO-UNT vs. WT-UNT DEG profile with that of Het-*AKAP11*-KO-Li vs. WT-UNT. Similarly, we then utilized the H3K27ac mark to determine the Li-rescued and -induced differential peaks and if such peaks were associated with the Li-rescued and -induced DEGs, respectively, in the same direction.

Among the transcriptomic effects of Li in the Het-*AKAP11*-KO iPSC-derived neuronal culture (Het-*AKAP11*-KO-Li vs. Het-*AKAP11*-KO-UNT), we found the downregulation of *GRIN2A*, which encodes the GluN2A subunit of NMDA receptors. *GRIN2A* has been reported to be associated with BD in GWAS^33,41^ and is a high-confidence LoF risk gene for SCZ^34,42,43^. Furthermore, it has been detected as a candidate gene in a GWAS of BD Li response^44^, suggesting a potential link between *GRIN2A* and Li treatment response in BD patients^44^. The encoded protein is involved in excitatory neurotransmission and synaptic plasticity^45,46^, and its expression regulation has been proposed as one of the therapeutic effects of Li in BD^47^. Interestingly, we showed that Li affected the gene expression profile of Het-*AKAP11*-KO distinctly from the WT model, with *GRIN2A* expression dysregulation being specific to Li treatment in Het-*AKAP11*-KO. These findings suggest that Li’s therapeutic effects in BD may be partly mediated by its ability to downregulate *GRIN2A*. This modulation may influence glutamatergic neurotransmission, which is reported to be dysregulated in BD^48,49^. Functional studies to investigate the specific role of identified genes such as *GRIN2A* in neurotransmission, in the presence and absence of Li in Het-*AKAP11*-KO model, represent an important direction for future research.

The absence of overlap between differential H3K27ac peaks and differential gene expression in the *AKAP11*-KO-Li vs. *AKAP11*-KO-UNT comparison is likely due to the subtle nature of Li’s effects on the chromatin (H3K27ac) landscape within this model. In contrast, clearer patterns of correspondence emerged when examining Li-rescued and Li-induced changes in H3K27ac peaks and gene expression relative to WT, with concordant directionality. These findings support a role for Li in enhancer-mediated transcriptional modulation. Specifically, they suggest that Li may contribute to the rescue of a subset of gene expression changes driven by AKAP11 haploinsufficiency, as well as the induction of new transcriptional alterations through the modulation of H3K27ac levels at intronic and intergenic regions, thereby influencing enhancer activity. The lack of correlation between differential H3K27ac peaks and DEGs in the direct *AKAP11*-KO-Li vs. *AKAP11*-KO-UNT comparison—where H3K27ac peaks were associated to the nearest TSS within 100 kb—also raises the possibility that Li-dependent enhancer changes within this model may influence more distal target genes, consistent with the well-established ability of enhancers to act over long genomic distances^50^. Therefore, the absence of DEG-H3K27ac overlap in this comparison (*AKAP11*-KO-Li vs. *AKAP11*-KO-UN) does not preclude the functional significance of these differential peaks, but rather underscores the complexity of enhancer-gene interactions and the need to incorporate three-dimensional genome architecture in future studies to investigate such long-range regulatory relationships.

The molecular functions related to the structural constituent of ribosomes and transcription regulation following chronic Li treatment in Het-*AKAP11*-KO were previously shown to be downregulated as a result of heterozygous knockout of *AKAP11*^25^. This downregulation also aligned with certain synapse proteomics GSEA findings from post-mortem dorsolateral prefrontal cortex samples of BD and SCZ patients compared to controls^30^, suggesting that Li may target and reverse key pathways commonly disrupted in Het-*AKAP11*-KO iPSC-derived neuronal cultures and in post-mortem brain tissues from BD and SCZ patients. Firstly, it is notable that there is a growing interest in studying the role of ribosome dysregulation induced by stress as a potential mechanism for neuropsychiatric disorders, particularly mood disorders^51–53^. Aberrant protein translation, possibly linked to decreased neurotransmission and synaptic plasticity^54,55^, is associated with the pathophysiology of SCZ^54,55^. In addition, dysfunctional stress response in the endoplasmic reticulum, which hosts a portion of the cytoplasmic ribosomes, is associated with BD^56,57^. Both in cell and animal models, Li has been shown to reverse stress-induced aberrant protein synthesis by decreasing the phosphorylation of a key modulator of mRNA translation eukaryotic elongation factor-2 (eEF2) at its inhibitory site, thereby increasing its activity, even under stress conditions such as nutrient and serum deprivation^58^. Secondly, mitochondrial abnormalities have been linked to major psychiatric disorders^59,60^. Mitochondria are crucial for neuronal functions, particularly in energy production, cellular respiration, and calcium signaling^61^. Evidence suggests that Li enhances mitochondrial respiration, possibly by influencing the mitochondrial oxidative phosphorylation (OXPHOS) pathway^61^. Consequently, the upregulation of mitochondrial ribosomal protein genes within the “structural constituent of ribosome” function observed after chronic Li treatment in the Het-*AKAP11*-KO model, may contribute to improved mitochondrial respiration. Lastly, Li can enhance the transcription and DNA binding through different paths. One possibility is via targeting the CREB (cAMP-response-element-binding protein)-co-activator TORC (transducer of regulated CREB) followed by enhancing CREB-directed gene transcription^62^. Other paths might involve Li targeting GSK3β, inositol mono phosphatase (IMPase), and/or inositol polyphosphate-1-phosphatase (IPPase)^63,64^ and affecting the downstream signaling and gene expression regulation.

Het-*AKAP11*-KO-Li vs. WT-UNT has a smaller number of DEGs than Het-*AKAP11*-KO-UNT vs. WT-UNT, under the same p.adj value and log2FC cutoffs, indicating that Li rescues some of the gene expression changes associated with the heterozygous knockout of *AKAP11*, bringing the Het-*AKAP11*-KO gene expression profile closer to that of the WT. This suggests a possible partial therapeutic or normalizing effect of Li on the Het-*AKAP11*-KO model. Chronic Li treatment of Het-*AKAP11*-KO rescued genes involved in various key roles, particularly glycosaminoglycan binding. Glycosaminoglycans are linear polysaccharides that exist on the cell surface in the ECM and covalently bind to core proteins to form proteoglycans^65^. Proteoglycans are one of the main components of the ECM. The Li-rescued genes associated with glycosaminoglycans binding function (*VTN, SMOC2, FGFBP1, HAPLN1, EGFLAM*) are primarily involved in the ECM organization, cell adhesion, and migration. The Li-induced DEGs are also involved in the function and structure of the ECM, signaling receptor regulator activity, calcium channel activity, and metal ion transmembrane transporter activity. Aberrant gene expression in ECM-related genes is noted in both Li-induced and Li-rescued genes, implying that Li’s therapeutic effects may involve complex regulation of ECM processes, which are crucial for neuronal activity, neuroplasticity, and cellular homeostasis^66^. Interestingly, alterations in ECM, such as degradation, overproduction, and altered composition have been reported in several neuropsychiatric disorders including BD, depression, and SCZ^66–68^.

We noted that the Li-responsive DEGs in Het-*AKAP11*-KO vs. WT, including both Li-rescued and Li-induced DEGs, were enriched in TCF4, LEF1, and PTX2 transcription factor binding sites. This suggests that one of the mechanisms by which Li may alter the expression of these genes is via its action on the GSK3β/β-catenin signaling pathway, which operates upstream of these transcription factors and is a well-established target of Li^21^. Western blot analysis of phosphorylated phGSK3β S9 revealed a significant decrease in phosphorylation levels in Het-*AKAP11*-KO neuronal cultures compared to WT counterparts. This finding is consistent with the reported role of AKAP11-dependent PKA activity in phosphorylating GSK3β at Ser9, a known direct target of this complex^15,19^. Heterozygous loss of AKAP11 function appears to impair PKA-mediated phosphorylation of GSK3β, leading to reduced inhibitory phosphorylation at this site. While this suggests impaired PKA-mediated regulation of GSK3β due to AKAP11 haploinsufficiency, it is important to consider that other kinases, such as Akt, protein kinase C (PKC), and p90RSK, can also phosphorylate GSK3β at Ser9 independently of AKAP11-anchored PKA activity. Despite these potential compensatory mechanisms, the considerable reduction in phosphorylation observed indicates that AKAP11 heterozygous loss has a notable impact on GSK3β activity regulation. Furthermore, we found that chronic Li treatment significantly increased pGSK3β Ser9 levels in Het-*AKAP11*-KO neurons, supporting Li’s known role in inhibiting GSK3β via enhancing Ser9 phosphorylation^31,32^. These findings suggest that Li’s inhibitory effect on GSK3β may contribute both to the partial rescue of transcriptomic alterations resulted from AKAP11 haploinsufficiency and to the induction of novel gene expression changes. While these rescuing and inducing effects may not act exclusively through pGSK3β pathways, our data suggests its contribution to Li’s broader molecular effects. Using iPSC-derived neurons from additional healthy individuals to generate more Het-*AKAP11*-KO isogenic lines, along with increasing the overall sample size of both KO and WT groups, would further enhance the statistical power and robustness of these findings.

Lastly, we showed that chronic Li treatment of Het-*AKAP11*-KO rescues a subset of DEGs resulting from heterozygous knockout of *AKAP11* and induces DEGs previously absent in Het-*AKAP11*-KO vs. WT iPSC-derived neurons. This effect partially occurs via Li’s impact on intergenic and intronic H3K27ac peaks, which mark active enhancers^27–29^, driving changes in the same direction as observed in gene expression. These particularly include non-coding RNA genes (lncRNAs and miRNAs), and coding genes involved in MAPK signaling cascades (e.g., *FGFR3*, *DUSP1*). *DUSP1* encodes MAP kinase phosphatase 1 (MKP1) which belongs to a family of proteins that dephosphorylate both threonine and tyrosine residues, acting as a negative regulator of the MAPK cascade^69^—a significant signaling pathway involved in neuronal plasticity, function, and survival^70,71^. Elevated *DUSP1* mRNA expression is associated with the pathophysiology of depressive behavior^72,73^. More specifically, elevated mRNA levels of *DUSP1* have been observed in the hippocampus of both patients with depression and stressed rats^72^. Overall, our findings suggest that Li can modify a subset of the gene expression changes that occur because of heterozygous knockout of *AKAP11* by altering H3K27ac levels at specific intergenic and intronic enhancers, thereby influencing enhancer activity and transcription factor binding affinity.

Several mood stabilizers used in the treatment of BD have been shown to modulate HDACs. For example, valproic acid acts as an HDAC inhibitor^74^ and lamotrigine has been reported to increase histone acetylation levels in vitro^75^. Notably, Li has been found to reduce HDAC1 protein expression without affecting its mRNA levels^76^. Although the precise mechanism remains unclear, evidence suggests that Li may interfere with the association of HDAC1 mRNA with polyribosomes, thereby impairing its translation^76^. In addition, chronic Li treatment in mice has been shown to significantly increase global histone H3 acetylation in specific brain regions^77,78^. These findings suggest that Li may modulate the aberrant enhancer-associated H3K27ac landscape observed in the Het-*AKAP11*-KO neuronal model, potentially through direct or indirect effects on HDACs such as HDAC1, although the exact mechanisms is not yet understood. Further research is needed to determine the involvement of HDACs in Li-associated H3K27ac changes in this model.

Due to the rarity of *AKAP11* LoF variants and the lack of available patient-derived iPSCs with confirmed *AKAP11* LoF mutations, we were unable to perform patient-specific comparisons. While our model provides valuable biological insights, future studies using iPSC-derived neurons from patients carrying *AKAP11* LoF variants, particularly those characterized as Li responders or non-responders, would enable more clinically relevant validation and offer deeper insight into the pathways identified and their connection to Li response in *AKAP11*-mutant cases.

Altogether, our findings enhance the understanding of how Li may exert therapeutic effects, particularly in individuals with BD and/or SCZ who carry *AKAP11* LoF variants. Additionally, Li may also benefit BD and SCZ patients who do not have this specific rare variant but exhibit alterations of similar pathways resulting from the heterozygous knockout of *AKAP11*, such as those related to ribosomes, proteoglycans, and ECM dynamics. These pathways were shown here to be influenced by Li and may be also critical to the phenotype. Overall, these findings highlight the complexity of gene expression regulation in response to Li treatment and emphasize that a specific genetic variation, such as heterozygous LoF mutations in *AKAP11*, may affect therapeutic outcomes. Although Li is not generally recommended for SCZ, this study opens the door to exploring its efficacy in a subset of patients based on their genetic background. Further research involving larger sample sizes, iPSC-derived neuronal cultures from BD and SCZ patients carrying *AKAP11* LoF variants, and functional studies is needed to advance our understanding of heterozygous *AKAP11* knockout in human cellular and animal models, as well as its associated response to Li in the context of BD and SCZ.

## Materials / Subjects and Methods

### Ethics statement

The cell line and protocols in the present study were used in accordance with guidelines approved by the institutional human ethics committee guidelines and the Nova Scotia Health Authority Research Ethics Board (REB #1020604) and the McGill University Health Centre (MUHC) Research Ethics Board (local REB #IRB00010120).

### RNP-CRISPR-Cas9 heterozygous knockout of *AKAP11* in induced pluripotent stem cells

The iPSCs used in this study were reprogrammed from the blood sample of a consenting individual (informed consent was obtained) as described in detail in our previous publication by Stern et al.^79^. Briefly, Epstein–Barr virus (EBV)-immortalized B-lymphocytes were reprogrammed using the Yamanaka episomal vector system^80^, consisting of three sets of episomal plasmids expressing reprograming factors: pCXLE-hOCT3/4-shp53, pCXLE-hSK (SOX2, KLF4) and pCXLE-hUL (L-MYC, LIN28)^79^. The protocol for the generation of Het-*AKAP11*-KOs with the RNP-CRISPR-Cas9 technique and the validation of the KOs using RT-qPCR and immunoblotting are described in our previous work^25^. The same WT replicates (SBP009, male, age 53, Caucasian; Table 1) and Het-*AKAP11*-KO clones (Het-*AKAP11*-KO-Clone 21: 4 bp insertion CAGT, Het-*AKAP11*-KO-Clone 16: 11bp deletion ACACCTTGGAC, Het-*AKAP11*-KO-Clone 20: 10 bp deletion GGTACACCTT) as our previous study^25^ are utilized here. Het-*AKAP11*-KO*s*: 3 clones each carrying a different frameshift mutation, all derived from SBP009; WT: 3 SBP009 replicates, each with a different passage number, cultured and differentiated to neurons separately.

**Table 1.** iPSC Sample Information.

| Sample | Source | Reprogramming | Identifier | Donor information |
| --- | --- | --- | --- | --- |
| Induced Pluripotent Stem Cell (iPSC) | Human lymphocytes | Reprogramming into iPSCs was performed at Sanford Burnham, as briefly described above and previously detailed by Stern et al.<sup>79</sup> | SBP009 | Control, Unaffected individual, Male, 51, Caucasian |

### iPSC and NPC differentiation and maintenance and neuronal cell culture

All iPSCs used in the present study were verified as free from mycoplasma contamination using Mycoplasma Detection Assays according to the manufacturer’s instructions (Thermo Fisher Scientific, Cat. #A55124). iPSCs were maintained on a Matrigel (Corning, Cat. #08-774-552) coated plate with mTeSR™1 medium (STEMCELL Technologies, Cat. #85870) along with Y-27632 (ROCK inhibitor, STEMCELL Technologies, Cat. # 72307) after each passage. During the course of induction of iPSC to NPCs (20 days), STEMdiff™ SMADi Neural Induction Kit (STEMCELL Technologies, Cat. #08581) was used. The method for induction of iPSC to NPCs was based on the STEMCELL™ neural induction EB-based method and was performed following the manufacturer’s protocol. On Day 0, a single-cell suspension of iPSC was generated using Gentle Cell Dissociation Reagent (STEMCELL Technologies, Cat. # 100-0485), and 10,000 cells per microwell were seeded in an ultralow attachment 96-well plate to generate Embryoid bodies (EBs), cultured in STEMdiff™ Neural Induction Medium + SMADi (STEMdiff™ SMADi Neural Induction Kit, Cat. # 08582) + 10 μM Y-27632. From Day 1 to Day 4, a daily partial (3/4) medium change was performed using STEMdiff™ Neural Induction Medium + SMADi. On Day 5, EBs were harvested using a wide-bore 1ml serological pipettes and 40μm strainer and transferred to a single well of a 6-well plate coated with Poly-L-ornithine hydrobromide (PLO) and laminin. A daily full medium change was performed from Day 6 to Day 11. After the neural induction efficiency was determined higher than 75%, neural rosettes were manually selected using STEMdiff™ Neural Rosette Selection Reagent (STEMCELL Technologies, Cat. # 05832) on Day 12 and replated onto a single well of PLO/Laminin coated 6-well plate. With continuous daily full medium change, selected rosette-containing clusters attached, and NPC outgrowths formed a monolayer between the clusters. The NPCs were ready for passage 1 when cultures were approximately 80-90% confluent (typically on Day 19). NPCs were maintained and expanded (7-10 days) using DMEM/F12 with Glutamax (ThermoFisher, Cat. #10565042), N2 1x (ThermoFisher, Cat. # 17502001), B27 1x (ThermoFisher, Cat. #17504001), FGF2 20 ng/ml (PEPROTECH, Cat. #100-18B), and EGF 20 ng/ml (Gibco, Cat. # AF-100-15-1MG). For the final differentiation of NPCs into neurons, NPCs were detached using Accutase (STEMCELL Technologies, Cat. #7922) and then differentiated onto PLO/laminin-coated plates with final differentiation media of BrainPhys™ Neuronal Medium (STEMCELL Technologies, Cat. #05790) supplemented with Glutamax 1x (ThermoFisher, Cat. #35050061), N2 1x (ThermoFisher, Cat. #17502001), B27 1x (ThermoFisher, Cat. #17504001), ascorbic acid 200nM (STEMCELL Technologies, Cat. #72132), cyclic AMP 500 μg/ml (Sigma-Aldrich, Cat #A9501-1G), brain-derived neurotrophic factor 20 ng/ml (BDNF, GIBCO, Cat. #PHC7074), Wnt3a 10 ng/ml (R&D Systems, Cat. #5036-WN-500), and laminin 1 μg/ml (Gibco, Cat. #23017015) for 14 days (medium change of 3 times per week). Subsequently, the cells were maintained in STEMdiff™ Forebrain Neuron Maturation Kit for 11 days (STEMCELL Technologies, Cat. #08605) in the same dishware with media change occurring every 4 days. At this stage and in parallel, a subset of the Het-*AKAP11*-KO dishes was treated with 1.5 mM Li included in the STEMdiff™ Forebrain Neuron Maturation Kit media. They were also maintained in the Li-containing media for 11 days, to mimic chronic Li treatment, with media change every 4 days. The culture composition was previously shown to be predominantly neuronal cells^25^. Other cell types, particularly glial cells, were present in lower proportions, establishing a mixed neuronal-glial culture, which we here refer to as a neuronal culture due to the dominance of neurons^25^.

### RNA-seq library Preparation and Sequencing

RNA was extracted from bulk neuronal culture 25 days post-differentiation using RNeasy Mini Kit (Cat. #74104). Library preparation from total RNA was performed using NEB rRNA-depleted (HMR) stranded library preparation kit according to the manufacturer’s instructions and sequencing was carried out using Illumina NovaSeq 6000 (100 bp paired-end).

### Immunoblotting

Cells were homogenized in lysis buffer (10LmM Tris–HCl pH7.5, 10LmM EDTA, 150LmM NaCl, 1% Triton X100, 0.1% SDS) in the presence of a mammalian protease inhibitor cocktail (Sigma, Cat #P8340) and phosphoSTOP (Sigma, Cat. #4906845001) to protect proteins from dephosphorylation. Protein extracts (30μg) were resolved by 10% Bis-tris gels, transferred onto PVDF membrane (Millipore IPFL00010), and immunoblotted with the primary antibody: rabbit polyclonal phospho-GSK-3 (Ser9; 1:1000, Cell Signaling Technology, Cat# 9336, RRID: AB_331405). Standard GAPDH loading controls were included using a mouse monoclonal anti-GAPDH antibody (1:2000, Invitrogen, Cat. #MA5-15738, RRID: AB_10977387). Subsequently, the membrane was revealed using the appropriate LI-COR fluorophore-conjugated secondary antibodies. Images were acquired using a LI-COR Odyssey Infrared image system. Fluorescence intensity values for each protein of interest were normalized to GAPDH signal from the same gel. Band quantification was carried out using Fiji software. Full-size blots for cropped gels can be found in the supplementary material. The two-way ANOVA followed by Tukey’s multiple comparison test was performed using GraphPad Prism version 10.2.3 (403) for Windows, GraphPad Software; www.graphpad.com.

### Cross-linking, ChIP-seq library preparation, and sequencing

About 10 million neuronal cells per sample (3 Het-*AKAP11*-KO clones and 3 WT replicates) were grown and directly crosslinked on the plate with 1% formaldehyde (Sigma) for 10 minutes at room temperature and the reaction was stopped using 125nM Glycine for 5 minutes. Fixed cell preparations were washed with ice-cold PBS, scraped off the plate, pelleted, washed twice again in ice-cold PBS, and the flash-frozen pellets were stored at −80°C. Thawed pellets were resuspended in 500ul cell lysis buffer (5 mM PIPES-pH 8.5, 85 mM KCl, 1% (v/v) IGEPAL CA-630, 50 mM NaF, 1 mM PMSF, 1 mM Phenylarsine Oxide, 5 mM Sodium Orthovanadate, EDTA-free Protease Inhibitor tablet) and incubated 30 minutes on ice. Samples were centrifugated and pellets resuspended in 500ul of nuclei lysis buffer (50 mM Tris-HCl pH 8.0, 10 mM EDTA, 1% (w/v) SDS, 50 mM NaF, 1 mM PMSF, 1 mM Phenylarsine Oxide, 5 mM Sodium Orthovanadate and EDTA-free protease inhibitor tablet) and incubated 30 minutes on ice. Sonication of lysed nuclei was performed on a BioRuptor UCD-300 at max intensity for 45 cycles, 10 s on 20 s off, centrifuged every 15 cycles, and chilled by a 4°C water cooler. Samples were checked for sonication efficiency using the criteria of 150–500bp by gel electrophoresis of a reversed cross-linked and purified aliquot. After the sonication, the chromatin was diluted to reduce the SDS level to 0.1% and concentrated using Nanosep 10k OMEGA (Pall). ChIP reaction for histone modifications was performed on a Diagenode SX-8G IP-Star Compact using Diagenode automated Ideal ChIP-seq Kit for Histones. Dynabeads Protein A (Invitrogen) were washed, then incubated with specific antibodies (rabbit polyclonal anti-H3K27ac Diagenode Cat. #C15410196, RRID: AB_2637079), 1 million cells of sonicated cell lysate, and protease inhibitors for 10 hr, followed by 20 min wash cycle using the provided wash buffers (Diagenode Immunoprecipitation Buffers, iDeal ChIP-seq kit for Histone). Reverse cross-linking took place on a heat block at 65°C for 4 hr. ChIP samples were then treated with 2ul RNase Cocktail at 65°C for 30 min followed by 2ul Proteinase K at 65°C for 30 min. Samples were then purified with QIAGEN MinElute PCR purification kit (QIAGEN) as per manufacturers’ protocol. In parallel, input samples (chromatin from about 50,000 cells) were reverse crosslinked, and DNA was isolated following the same protocol. Library preparation was carried out using Kapa Hyper Prep library preparation reagents (Kapa Hyper Prep kit, Roche 07962363001) following the manufacturer’s protocol. ChIP libraries were sequenced using Illumina NovaSeq 6000 at 100bp paired-end reads.

### RNA-seq data processing and analysis

Adaptor sequences and low-quality score bases (Phred score < 30) were first trimmed using Trimmomatic^81^. The resulting reads were aligned to the GRCh38 human reference genome assembly, using STAR^82^. Read counts were obtained using HTSeq^83^ with parameters-m intersection-nonempty-stranded=reverse, using version Ensembl112 GTF annotation. For all downstream analyses, we excluded lowly-expressed genes with an average read count lower than 20 across all samples and unannotated genes (with ID names starting with ENSG), resulting in 19,568 genes. Raw counts were normalized using edgeR’s TMM algorithm^84^ and were then transformed to log2-counts per million (logCPM) using the voom function implemented in the limma R package^85^. To assess differences in gene expression levels between the different conditions (3 KO clones versus 3 WT replicates), we fitted a linear model using limma’s lmfit function with parameter method=“robust”. For the Li-plus vs Li-minus analyses, we used a paired statistical test to account for the sample matchings between groups. Nominal p-values were corrected for multiple testing using the Benjamini-Hochberg method. The Z-score heatmap was constructed using the R package ComplexHeatmap^86^; KO vs WT analysis: P-adj (adjusted p-value) < 0.05 & |log2FC| > 0.25. Gene set enrichment analysis (GSEA)^36,39^ based on a pre-ranked gene list by t-statistic was performed using the R package fgsea (http://bioconductor.org/packages/fgsea), P-adj < 0.05. Over-representation analysis (ORA) was performed using ShinyGO 0.77^40^ (http://bioinformatics.sdstate.edu/go). The background genes used for ORA were all the 19,568 genes that were considered expressed. FDR is calculated based on the nominal P-value from the hypergeometric test. Fold Enrichment is the percentage of genes in the input gene list belonging to a pathway divided by the corresponding percentage in the background. For visualization on the heatmaps, gene expression data was corrected for batch effects using ComBat^87^. For track visualizations, Bigwig coverage files were created with the HOMER makeUCSCfile command and bedGraphToBigWig utility from UCSC. Data were normalized so that each value represents the read count per base pair per 10 million reads. UCSC Genome Browser (http://genome.ucsc.edu/) was implemented for track visualization. Significantly differentially expressed genes in KO-Li vs KO-UNT comparison were obtained using the cutoffs P.adj < 0.05 and |log2FC| > 0.25. Li-induced/rescued genes were obtained using the criteria below:

- Significantly differentially expressed (P.adj < 0.05 and |log2FC| > 0.25) in one comparison but not the other (i.e., in either Het-*AKAP11*-KO-UNT vs. WT-UNT or Het-*AKAP11*-KO-Li vs. WT-UNT)
- |diff_log2FC| > 0.6

### ChIP-seq data processing and analysis

ChIP-seq reads were first trimmed for adapter sequences and low-quality score bases using Trimmomatic^81^. The resulting reads were mapped to the human reference genome (GRCh38) using BWA-MEM^88^ in paired-end mode at default parameters. Only reads that had a unique alignment (mapping quality > 20) were retained and PCR duplicates were marked using Picard tools (http://broadinstitute.github.io/picard/). Peaks were called using MACS2^89^. To detect changes in histone modifications, a consensus peak set for H3K27ac was first generated by merging ChIP-seq peaks across samples using bedtools merge (https://bedtools.readthedocs.io/). A peak must be present in at least one sample in either condition. Read counts were obtained within these genomic regions using HOMER. Differential peak analysis was performed using the R package limma^85^. Nominal p-values were corrected for multiple testing using the Benjamini-Hochberg method. Peaks were associated with the nearest TSS of genes using the annotatePeaks command from HOMER software suite^90^. Enrichment ORA of genes assigned to differential peaks was performed using ShinyGO 0.77^40^ (http://bioinformatics.sdstate.edu/go). The background genes used for ORA were all the 19,568 genes that were considered expressed. FDR is calculated based on the nominal P-value from the hypergeometric test. Fold Enrichment is the percentage of genes in the input gene list belonging to a pathway divided by the corresponding percentage in the background. Bigwig coverage files were created with the HOMER makeUCSCfile command and bedGraphToBigWig utility from UCSC. Data were normalized so that each value represents the read count per base pair per 10 million reads. UCSC Genome Browser (http://genome.ucsc.edu/) was implemented for track visualization. Significantly differentially marked peaks in KO-Li vs KO-UNT comparison were obtained using the cutoffs P.adj < 0.05 and |log2FC| > 0.25. Li-induced/rescued peaks were obtained using the criteria below:

- Significantly differentially marked (P.adj < 0.05 and |log2FC| > 0.25) in one comparison but not the other (i.e., in either Het-*AKAP11*-KO-UNT vs. WT-UNT or Het-*AKAP11*-KO-Li vs. WT-UNT)
- |diff_log2FC| > 0.3

## Acknowledgments

The work in GAR’s lab is supported by the ERA-PerMed in partnership with FRQS (JTC2018, 280240). BC’s team is supported by ERAPerMed PLOT-BD, a grant from the Fondation Bettencourt Schueller, and a government grant managed by the Agence Nationale de la Recherche under the France 2030 program (ANR-22-EXPR-0013). We also acknowledge the financial support from the Brain & Behavior Research Foundation Young Investigator Award to AK (30822). NF is supported by the studentship award Fonds de Recherche du Québec – Santé (FRQS): https://doi.org/10.69777/320249. CL is supported by the CIHR Banting Fellowship. Computational analysis, data pre-processing, and adapted bioinformatics pipelines for analyses were performed by the Canadian Centre for Computational Genomics (C3G)-Montréal Node using infrastructure provided by Compute Canada and Calcul Quebec. ChIP-seq library preparation and sequencing were performed at the McGill Genome Center platform.

## Author contributions

NF conceived and designed the study, defined its scope, and conducted the laboratory experiments and the cell culture work. NF planned the analytical approach, and AP (Pacis) conducted the computational data analysis and generated most of the plots and heatmaps included in the figure panels. Results interpretation was performed by NF. YL and AK provided expert guidance and assistance on the maintenance and handling of iPSCs, NPCs, and neuronal cultures. AP (Pietrantonio) and DR assisted with laboratory work related to the western blotting experiments. MA, PAD, GAR, BC, CL, and AK provided valuable scientific insights throughout the study and revised the manuscript. NF wrote the manuscript, and all authors reviewed the final version.

## Additional Information

### Data availability

Data are available at NCBI GEO under the accession numbers GSE290452 (RNA-seq data) and GSE290454 (ChIP-Seq data). All other relevant data supporting the key findings of this study are available within the article and its Supplementary Information files or from the corresponding authors upon request.

### Materials availability

This study did not generate new unique reagents.

### Lead contacts

Additional information and requests for resources and reagents should be directed to and will be fulfilled by the Lead Contacts Drs. Rouleau, Khayachi, and Chaumette.

