## Supplementary Information File for "Lithium partially rescues gene expression and enhancer activity from heterozygous knockout of *AKAP11* while inducing novel differential changes"

### Supplementary Figures:

**Supplementary Fig. S1 (Related to Fig. 1).** Transcriptomic effects of Li within the Het-*AKAP11*-KO and WT iPSC-derived neuronal models.

**Supplementary Fig. S2 (Related to Fig. 2).** Li's role in rescuing some of the DEGs obtained from Het-*AKAP11*-KO-UNT vs. WT-UNT and inducing new DEGs previously absent in Het-*AKAP11*-KO-UNT vs. WT-UNT.

**Supplementary Fig. S3 (Related to Fig. 3).** Differential intergenic and intronic H3K27ac peaks from Het-*AKAP11*-KO-Li vs. Het-*AKAP11*-KO-UNT comparison.

### Supplementary Tables:

Supplementary Tables S1-S10 captions, corresponding to the large data tables provided in Excel format separately.

### Supplementary Figures

Supplementary Fig. S1

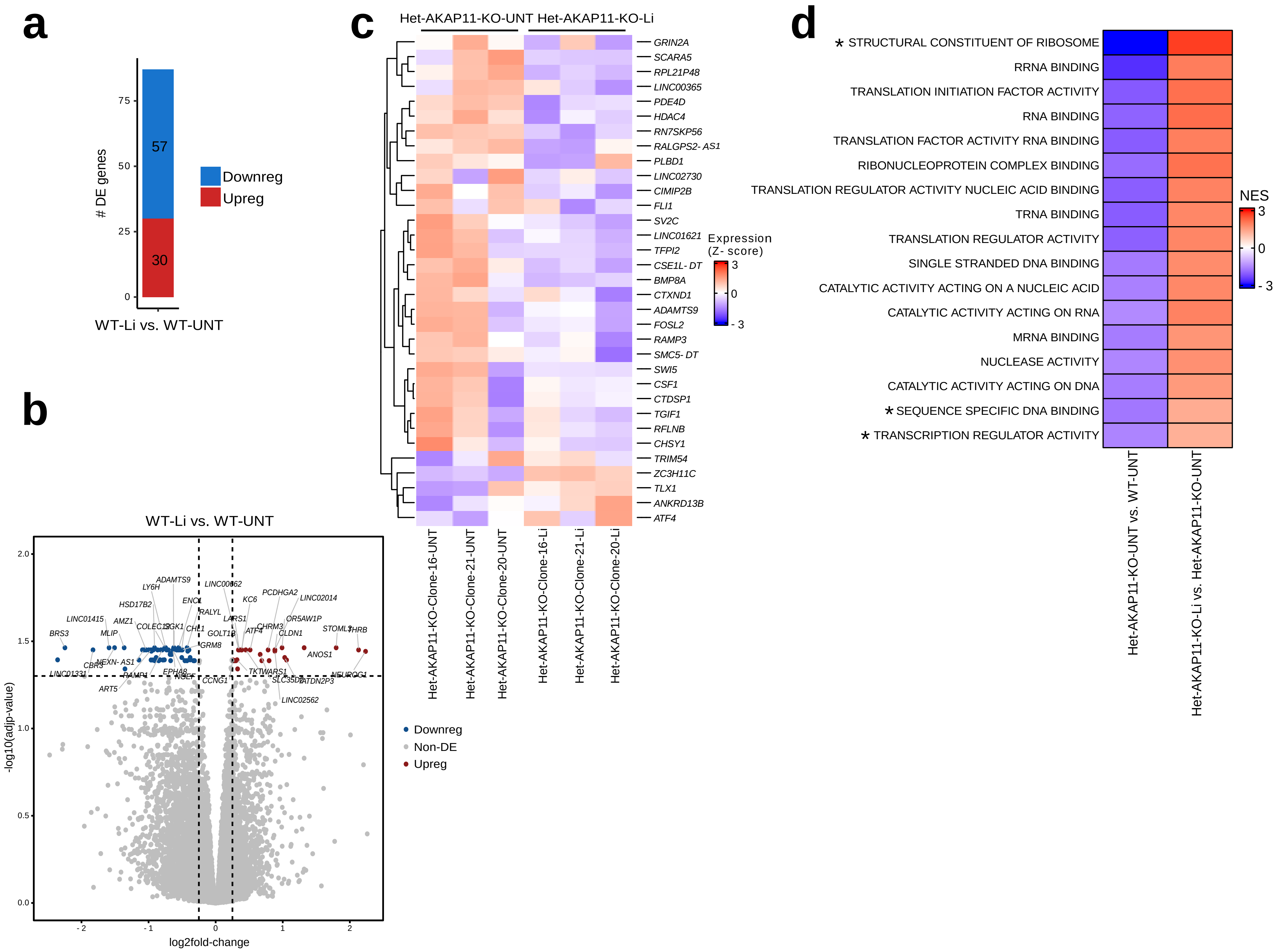

**Supplementary Fig. S1 (Related to Fig. 1). Transcriptomic effects of Li within the Het-*AKAP11*-KO and WT iPSC-derived neuronal models.** (**a**) Number of up and downregulated DEGs in WT-Li vs. WT-UNT; p.adj < 0.05 and |log2FC| > 0.25. Red: upregulated, blue: downregulated. (**b**) Volcano plot of differentially expressed genes from WT-Li vs. WT-UNT comparison; p-value < 0.05 and |log2FC| > 0.25. Red: upregulated, blue: downregulated, grey: nonDEG. (**c**) Hierarchical clustering heatmap of differentially expressed genes (rows) from each heterozygous Het-*AKAP11*-KO-Li clones vs. Het-*AKAP11*-KO-UNT reps. The expression levels across samples were standardized by the Z-Score method. Red shows the highest (3) and blue the lowest (-3) Z-score values. (**d**) Heatmap of normalized enrichment scores (NES) of the overlapping GSEA[36,39] GO[37,38] molecular functions between Het-*AKAP11*-KO-Li vs. Het-*AKAP11*-KO-UNT and Het-*AKAP11*-KO-UNT vs. WT-UNT from our previous study[25]. The functions marked with * are those that were previously shown to be also significantly downregulated (p.adj < 0.1) in the GSEA of synapse proteomics from post-mortem brain samples of BD and SCZ patients relative to controls[30] as well as in our iPSC-derived Het-*AKAP11*-KO neuronal culture[25].

Supplementary Fig. S2

a

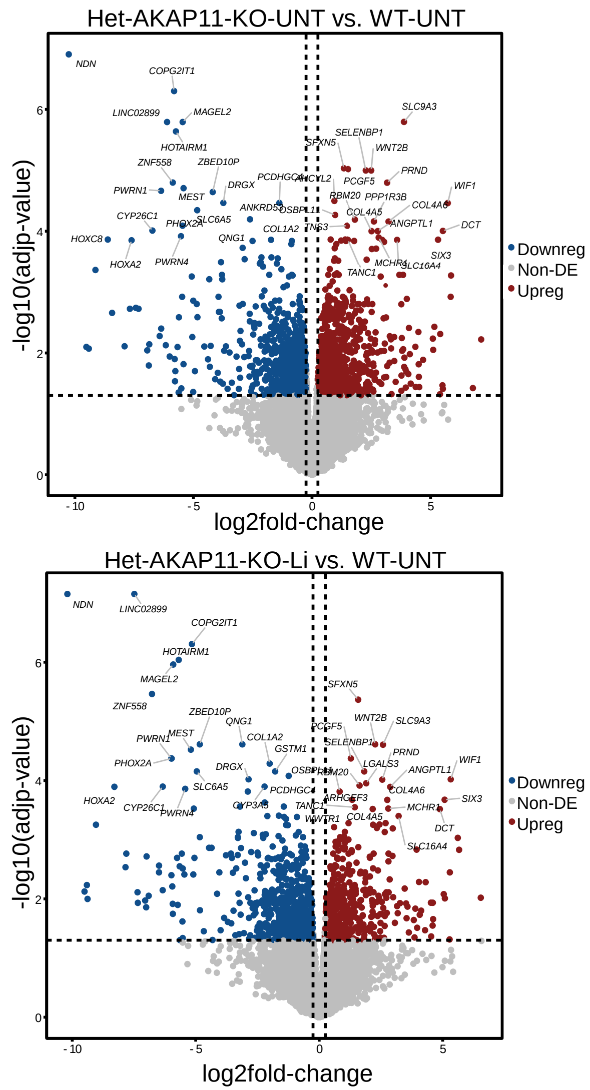


b

| Pathway | Enrichment FDR | nGenes | Pathway Genes | Fold Enrichment | Genes |
|---|---|---|---|---|---|
| Glycosaminoglycan binding, GO:0005539 | 5.86E-02 | 5 | 253 | 8.824 | VTN SMOC2 FGFBP1 HAPLN1 EGFLAM |
| Signaling receptor activity, GO:0038023 | 1.03E-01 | 7 | 1940 | 2.580 | FGFR3 GAL FZD10 RAMP3 CHRND GLRA1 SPHK1 |
| G protein-coupled receptor activity, GO:0004930 | 1.03E-01 | 4 | 1026 | 3.988 | GAL FZD10 RAMP3 SPHK1 |
| Growth factor binding, GO:0019838 | 1.03E-01 | 4 | 152 | 9.165 | FGFR3 DUSP1 FGFBP1 FGFBP2 |
| Ligand-gated ion channel activity, GO:0015276 | 1.03E-01 | 3 | 147 | 7.412 | CHRND GLRA1 RYR3 |
| Excitatory extracellular ligand-gated ion channel activity, GO:0005231 | 1.03E-01 | 2 | 33 | 21.855 | CHRND GLRA1 |
| Oxidoreductase activity, acting on single donors with incorporation of molecular oxygen, GO:0016701 | 1.03E-01 | 2 | 30 | 31.569 | BCO1 HPD |
| Fibroblast growth factor binding, GO:0017134 | 1.03E-01 | 2 | 23 | 27.059 | FGFR3 FGFBP1 |
| DNA-binding transcription factor activity, RNA polymerase II-specific, GO:0000981 | 1.47E-01 | 8 | 1446 | 2.044 | MEOX1 FOXC1 FOSL2 NKX3-2 IRX4 HOXD11 SHOX2 IRX3 |
| Transmembrane transporter activity, GO:0022857 | 1.98E-01 | 6 | 1248 | 1.935 | SLC7A8 ABCC3 SLC14A2 CHRND GLRA1 RYR3 |

c

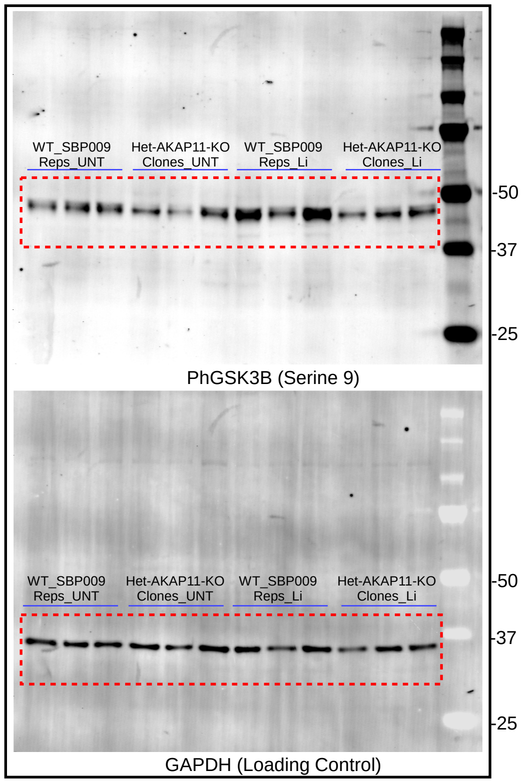

**Supplementary Fig. S2 (Related to Fig. 2). Li's role in rescuing some of the DEGs obtained from Het-*AKAP11*-KO-UNT vs. WT-UNT and inducing new DEGs previously absent in Het-*AKAP11*-KO-UNT vs. WT-UNT.** (**a**) Volcano plot of DEGs from Het-*AKAP11*-KO-UNT vs. WT-UNT (top) and Het-*AKAP11*-KO-Li vs. WT-UNT (bottom) comparisons; p-value < 0.05 and |log2FC| > 0.25. Red: upregulated, blue: downregulated, grey: non-DEG. (**b**) ORA of rescued DEGs following Li treatment of Het-*AKAP11*-KO (using all genes from Fig. 2b heatmap). GO[37,38] Molecular function database using ShinyGO 0.77[40]; Included in this table are FDR < 0.2. The yellow highlight indicates the most significant GO term. The background genes used were all the 19,568 genes that were considered expressed (see Methods). Shown here are the top 10 pathways, with redundancy removed. (**c**) Full immunoblot membranes including all 12 samples; top: phGSK3B (S9), bottom: GAPDH loading control (both from the same gel). Red rectangles highlight the sections shown in the cropped membranes in Fig. 2.

Supplementary Fig. S3

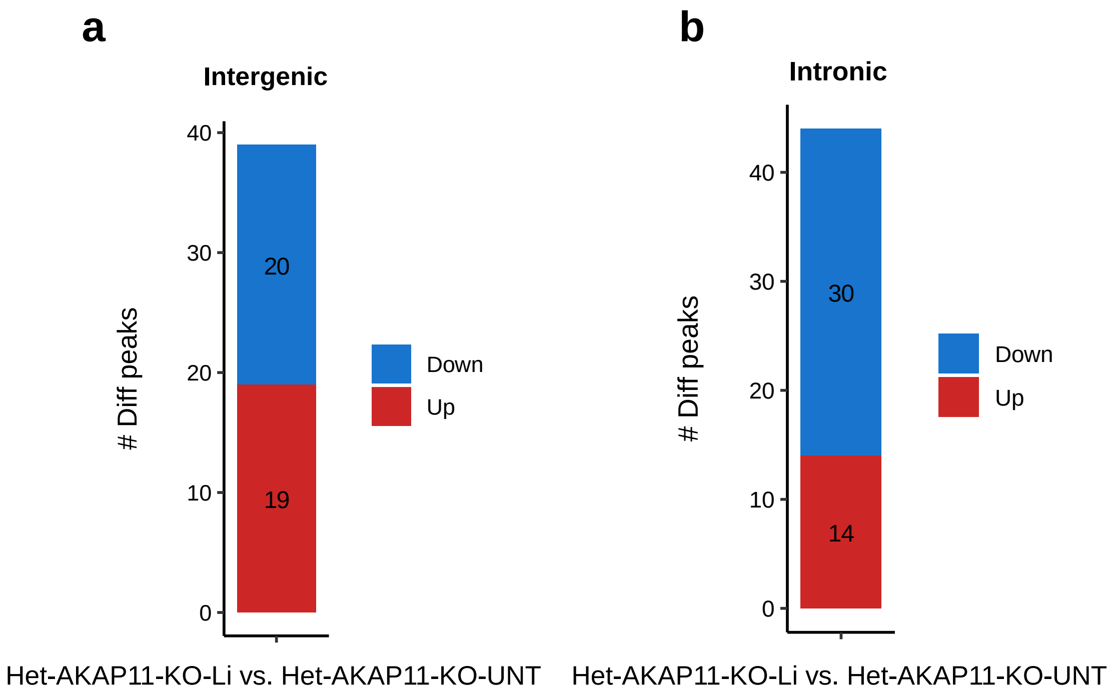

**Supplementary Fig. S3 (Related to Fig. 3). Differential intergenic and intronic H3K27ac peaks from Het-*AKAP11*-KO-Li vs. Het-*AKAP11*-KO-UNT comparison.** (**a**) **Differential intergenic peaks from** *AKAP11*-KO-Li vs. Het-*AKAP11*-KO-UNT comparison**;** p.adj < 0.05 and |log2FC| > 0.25; red: differentially elevated peaks, blue: differentially decreased peaks. (**b**) Differential intronic peaks from *AKAP11*-KO-Li vs. Het-*AKAP11*-KO-UNT comparison; P.adj < 0.05 and |log2FC| > 0.25; red: differentially elevated peaks, blue: differentially decreased peaks.

**Supplementary Tables Captions (**corresponding to the large data tables provided in Excel format separately**)**

**Supplementary Table S1.** DEG results of all the comparisons. We used DEGs with the cutoff of p-value < 0.05 and |log2FC| > 0.25 in this study.

**Supplementary Table S2.** GSEA[36,39] GO[37,38] molecular function of Het-*AKAP11*-KO-Li vs. Het-*AKAP11*-KO-UNT; P.adj<0.05.

**Supplementary Table S3.** Overlapping GSEA[36,39] GO[37,38] molecular functions between Het-*AKAP11*-KO-Li vs. Het-*AKAP11*-KO-UNTand Het-*AKAP11*-KO-UNT vs. WT-UNT from our previous study[25].

**Supplementary Table S4.** Li-rescued and Li-induced DEG list. This table shows those genes that are significantly differentially expressed (P.adj < 0.05 and |log2FC| > 0.25) in one comparison but not the other (in either Het-*AKAP11*-KO-UNT vs. WT-UNT or Het-*AKAP11*-KO-Li vs. WT-UNT); |diff_log2FC| > 0.6.

**Supplementary Table S5.** Transcription factor Target Genome Browser PWMs enrichment analysis generated using ShinyGo 0.77[40]. The table includes the top 20 most significant enriched motifs with FDR<0.1.

**Supplementary Table S6.** Immunoblot quantification table Het-*AKAP11*-KO-UNT, WT-UNT Het-*AKAP11*-KO-Li, WT-Li, determined using Fiji software.

**Supplementary Table S7.** Het-*AKAP11*-KO-Li vs. Het-*AKAP11*-KO-UNT intergenic and intronic differential H3K27ac peaks and associated genes within the distance of 100 kb from the TSS to the peak center along with their gene expression levels.

**Supplementary Table S8.** Differential (p.adj < 0.05 and |log2FC| > 0.25) intergenic and intronic H3K27ac peaks annotated results for Het-*AKAP11*-KO-UNT vs WT-UNT and Het-*AKAP11*-KO-Li vs. WT-UNT, along with all other comparisons.

**Supplementary Table S9.** Li-responsive differential intergenic and intronic H3K27ac peaks. Significantly differentially marked (P.adj < 0.05 and |log2FC| > 0.25) in one comparison but not the other (in either Het-*AKAP11*-KO-UNT vs. WT-UNT or Het-*AKAP11*-KO-Li vs. WT-UNT; |diff_log2FC| > 0.3). The H3K27ac peak-associated genes that overlapped with all Li-rescued or Li-induced DEGs are noted at the top. Yellow highlights indicate Li-rescued and Li-induced peaks that were linked to Li-rescued or Li-induced DEGs, respectively, with the same direction of change.

**Supplementary Table S10.** Over-representation analysis results (Curated.Reactome database) using the Li-recued and Li-induced intergenic and intronic H3K27ac-DEGs (genes listed in Fig. 3c, d). Generated with ShinyGo 0.77[40]. FDR<0.05, redundancy removed.
